# Pyrite formation from FeS and H_2_S is mediated by a novel type of microbial energy metabolism

**DOI:** 10.1101/396978

**Authors:** Joana Thiel, James Byrne, Andreas Kappler, Bernhard Schink, Michael Pester

## Abstract

The exergonic reaction of FeS with H_2_S to form FeS_2_ (pyrite) and H_2_ was postulated to have operated as an early form of energy metabolism on primordial Earth. Since the Archean, sedimentary pyrite formation played a major role in the global iron and sulfur cycles, with direct impact on the redox chemistry of the atmosphere. To date, pyrite formation was considered a purely geochemical reaction. Here, we present microbial enrichment cultures, which grew with FeS, H_2_S, and CO_2_ as their sole substrates to produce FeS_2_ and CH_4_. Cultures grew over periods of three to eight months to cell densities of up to 2–9×10^6^ cells mL^−1^. Transformation of FeS with H_2_S to FeS_2_ was followed by ^57^Fe Mössbauer spectroscopy and showed a clear biological temperature profile with maximum activity at 28°C and decreasing activities towards 4°C and 60°C. CH_4_ was formed concomitantly with FeS_2_ and exhibited the same temperature dependence. Addition of either penicillin or 2-bromoethanesulfonate inhibited both FeS_2_ and CH_4_ production, indicating a syntrophic coupling of pyrite formation to methanogenesis. This hypothesis was supported by a 16S rRNA gene-based phylogenetic analysis, which identified at least one archaeal and five bacterial species. The archaeon was closely related to the hydrogenotrophic methanogen *Methanospirillum stamsii* while the bacteria were most closely related to sulfate-reducing *Deltaproteobacteria*, as well as uncultured *Firmicutes* and *Actinobacteria*. We identified a novel type of microbial metabolism able to conserve energy from FeS transformation to FeS_2_, which may serve as a model for a postulated primordial iron-sulfur world.

**Significance statement:** Pyrite is the most abundant iron-sulfur mineral in sediments. Over geological times, its burial controlled oxygen levels in the atmosphere and sulfate concentrations in seawater. Its formation in sediments is so far considered a purely geochemical process that is at most indirectly supported by microbial activity. We show that lithotrophic microorganisms can directly transform FeS and H_2_S to FeS_2_ and use this exergonic reaction as a novel form of energy metabolism that is syntrophically coupled to methanogenesis. Our results provide insights into a syntrophic relationship that could sustain part of the deep biosphere and lend support to the iron-sulfur-world theory that postulated FeS transformation to FeS_2_ as a key energy-delivering reaction for life to emerge.

## Introduction

With an annual formation of at least 5 million tons, pyrite (FeS_2_) is the thermodynamically stable end product of iron compounds reacting with sulfide in reduced sediments, with the latter being produced mainly by microbial sulfate reduction. Consequently, pyrite is the most abundant iron-sulfur mineral on Earth’s surface (1). Over geological times, burial of pyrite was tightly intertwined with organic matter preservation in reduced sediments (2). These massive reservoirs of reduced sulfur and carbon are being counterbalanced by the photosynthetically produced oxygen in the Earth’s atmosphere (2). Despite this importance of pyrite for Earth’s iron, sulfur, and carbon cycles as well as Earth’s surface redox state, the mechanism of pyrite formation in natural environments is still being debated (1). Currently, two mechanisms are discussed to drive pyrite formation in sediments, which both preclude dissolution of precipitated iron(II) monosulfide (FeS) to an aqueous FeS intermediate. In the polysulfide pathway, FeS_aq_ is attacked by nucleophilic polysulfide to form FeS_2_ (equ. 1). Alternatively, pyrite may form from the reaction of FeS_aq_ with H_2_S (equ. 2), which is known as the H_2_S pathway or the Wächtershäuser reaction (1).

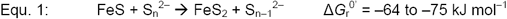

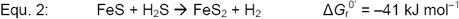

Using inorganic experimental systems, abiotic pyrite formation has been observed at temperatures of 25–125°C (e.g. 1, 3). However, already trace amounts of organic compounds containing aldehyde groups were reported to inhibit pyrite formation in such experiments (1, 4). Absence of stringent control for such compounds may explain why many other studies with abiotic systems did not observe pyrite formation at environmentally relevant temperatures (5-11). On the other hand, pyrite formation is known to take place in the presence of organic matter and especially alive microorganisms in sediments (5, 12). Indeed, pyrite formation could be observed as a side reaction in pure and enrichment cultures of heterotrophic sulfate-reducing and chemolithotrophic sulfur-dismutating bacteria, with the assumption that these microorganisms provide mainly H_2_S as a substrate for concomitant abiotic pyrite formation (13-15). In addition, a more complex involvement of microorganisms has been postulated that includes the support of nucleation of FeS minerals on bacterial cell surfaces (16). However, in all these studies biogenic pyrite formation has so far been understood only as an indirect abiotic process.

Pyrite formation according to equ. 2 provides reducing equivalents in the form of H_2_ that could be coupled to the reduction of CO_2_ to CH_4_ or more complex organic matter. Coupling of pyrite formation to methanogenesis has been proposed by Jørgensen and coworkers to be part of a cryptic sulfur cycle in deep marine sediments where it could support the enigmatic life forms of the deep biosphere (17). Coupling of this reaction to the synthesis of organic matter is the basis of the “iron-sulfur world” theory proposed by Wächtershäuser, by which pyrite formation is viewed as the central process that led to the transition from Fe-S surface-catalyzed synthesis of organic molecules to actual life on the primordial Earth (18-20). Here, we show for the first time that the reaction of FeS with H_2_S to form FeS_2_ can be used by microorganisms to conserve energy for lithotrophic growth if syntrophically coupled to methanogenesis.

## Results and Discussion

### Pyrite formation as a microbially catalyzed process

Mineral medium containing 5 mM FeS and 6 mM H_2_S as sole substrates and CO_2_/HCO_3_^−^ as carbon source was inoculated with digested sewage sludge, freshwater or marine sediment (Table S1) and incubated at 16°C or 28°C. Microbial activity was followed via methane formation, and transfers were made every three to eight months, typically when the methane content in the headspace approached a plateau. Of seven enrichments started, four exhibited methane formation for more than ten transfers. The most active enrichment culture J5, which was started from digested sewage sludge and incubated at 28°C, was characterized in more detail after more than 20 transfers. On average, the methane content reached 0.7 mmol per L of culture in J5. In contrast, no methane formation was observed in abiotic controls (Fig. 1*A*). This was mirrored in the turnover of total H_2_S: While in culture J5 total H_2_S decreased over time from approx. 6 mmol to 0.04–1.1 mmol per L of culture (Fig. S1), abiotic controls showed a much less pronounced decline of total H_2_S (3.7 mmol residual H_2_S per L). The observed decrease in the abiotic controls may be due to inorganic background reactions (see below).

**Figure 1.**
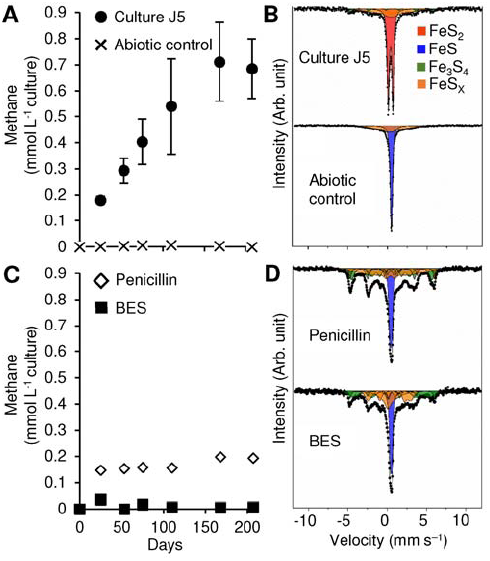
Time-resolved CH_4_ formation in comparison to iron-sulfur mineral composition after nearly seven months of incubation (207 days) in culture J5 as compared to abiotic controls and incubations of culture J5 with penicillin-G (1,000 U ml^−1^) or 2-bromoethanesulfonate (BES, 10 mM). (A) and (C) show the mean ± one standard deviation of CH_4_ measurements of three independent incubations. Standard deviations are often smaller than the actual symbol size. (B) and (D) show Mössbauer spectra corresponding to the last time point in the presented time series with FeS_2_ in red, FeS in blue, Fe_3_S_4_ in green, and intermediate FeS_x_ phases in orange.

Conversion of FeS solids was followed by ^57^Fe Mössbauer spectrometry. After nearly seven months of incubation, the Mössbauer spectrum of culture J5 was dominated by a FeS_2_ doublet (Fig. 1*B*), which corresponded to 53–63% of the iron-sulfur mineral phase (Table S2). In contrast, no evidence of a singlet peak corresponding to FeS was present. In addition, a poorly defined sextet feature was required to achieve a satisfactory fit of the Mössbauer spectrum. This poorly defined sextet is best described as a metastable phase, which we have termed FeS_x_ in accordance with the notation used by Wan et al. (21), and represented 31–39% of the remaining iron-sulfur minerals. Furthermore, a well-defined sextet with a hyperfine magnetic field of 32 T was required to fit the data, which corresponded well to greigite (Fe_3_S_4_) (22) and made up 7–8% of the remaining iron-sulfur minerals. Greigite is the sulfur isomorph of magnetite and was previously observed as an intermediate phase in the FeS conversion to pyrite in abiotic studies (23, 24).

FeS_2_ formation in culture J5 was confirmed by X-ray diffraction analysis, which recovered all major XRD reflections of pyrite in the obtained XRD pattern (Fig. 2*A*). Since no indication of a parallel formation of the dimorph marcasite was evident from the XRD pattern, the recovered FeS_2_ phase is referred to as pyrite from here on. Further support for pyrite formation in culture J5 was provided by scanning electron microscopy (SEM) coupled to energy-dispersive X-ray (EDX) spectroscopy. Here, µm-scale iron-sulfur crystals with a euhedral structure as typical of pyrite could be observed (Fig. 2*B*), which resembled the expected Fe:S ratio of 1:2 as revealed by EDX point measurements (Fig. S2).

**Figure 2.**
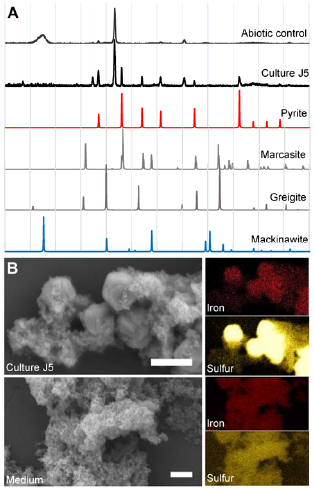
(A) Representative X-ray diffractograms of mineral precipitates formed in culture J5 and an abiotic control setup after 9 months of incubation (281 days). Diffraction data of the two FeS_2_ dimorphs pyrite and marcasite as well as of Fe_3_S_4_ (greigite) and FeS (mackinawite) are given as reference. (B) Scanning electron microscopy images of a nearly 7-months old (211 days) culture J5 in comparison to freshly prepared medium without inoculum. The scale bar represents 2 µm. Images to the right show the corresponding results from energy dispersive X-ray spectroscopy (EDX). Besides atoms from medium salts, iron and sulfur were the only elements discovered in the mineral phases.

In contrast to culture J5, the Mössbauer spectrum of the nearly seven-months-old abiotic control was dominated by a prominent FeS singlet peak (64% of iron-sulfur minerals). In addition, a poorly defined sextet corresponding to FeS_x_ was required to achieve a satisfactory fit of the obtained data (36% of iron-sulfur minerals) (Fig. 1, Table S2). The abiotic conversion of FeS to FeS_x_ most likely explains the observed decrease of total H_2_S in the abiotic control. Absence of pyrite formation in abiotic controls was further supported by the obtained XRD patterns (Fig. 2*A*). In addition, freshly prepared medium was also devoid of pyrite as evidenced by an overall disordered iron-sulfur mineral phase in SEM-EDX images without any euhedral crystals indicative of pyrite (Fig. 2*B*, Fig. S2).

To further support our hypothesis of biogenic pyrite formation in culture J5, we followed its iron-sulfur mineral transformation over a temperature gradient of 4–60°C. The maximum of pyrite formation was evident at 28°C. Incubation at lower (16°C) or higher (46°C) temperatures resulted in decreased pyrite formation, with no pyrite formation at 4°C and 60°C (Fig. 3*A* and 3*B*, Table S2). While FeS was completely transformed to pyrite, greigite, and various FeS_x_ phases at 16, 28, and 46°C, some residual FeS remained at 4°C and 60°C (Fig. 3*A*). The observed temperature profile of pyrite formation is typical of biologically catalyzed reactions centered on an optimum reaction temperature. In contrast, abiotic pyrite formation at temperatures of <100°C was shown to follow a sigmoidal temperature dependence (3).

**Figure 3.**
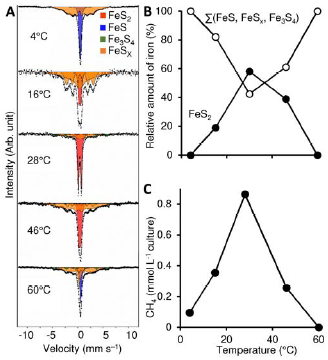
Temperature-dependent pyrite and methane formation in culture J5 after nearly 7 months of incubation (207 days). A) Mössbauer spectra showing the temperature-dependent iron-sulfur mineral composition (FeS_2_ in red, FeS in blue, Fe_3_S_4_ in green, and intermediate FeS_x_ phases in orange). B) Relative abundance of pyrite (FeS_2_) in comparison to all other measured iron-sulfur minerals plotted against temperature as the explanatory variable. Details are provided in Table S2. C) Average amount of methane (n=2) plotted against temperature as the explanatory variable.

### Microbial pyrite formation is a syntrophically coupled process

Methane formation closely followed the temperature-dependent activity profile of pyrite formation (Fig. 3*C*) suggesting metabolic coupling of both processes. This hypothesis was further explored in an inhibition experiment using either penicillin G as a generic inhibitor of bacterial cell wall synthesis or 2-bromoethanesulfonate (BES) as a specific inhibitor of hydrogenotrophic methanogenesis (25). BES inhibited both methane and pyrite formation completely (Fig. 1*C* and 1*D*). We interpret this as a direct coupling of biogenic pyrite formation to hydrogenotrophic methanogenesis. In support of this hypothesis, penicillin inhibited both pyrite and methane formation as well (Fig. 1*C* and 1*D*). Here, methane formation ceased after an initial production of 0.15 mmol per L of culture. The latter is explained by penicillin’s mode of action, which inhibits growth of bacteria but not their initial metabolic activity. Interestingly, a corresponding small amount of pyrite was not observed in this experiment but rather a partial FeS transformation to various FeS_x_ phases and greigite (Fig. 1*D*, Table S2). This indicates that methanogenesis could receive reducing equivalents also from these iron-sulfur mineral transformations. Further support for syntrophic coupling of pyrite formation to methanogenesis was provided by a third inhibition experiment in which penicillin addition was supplemented by 79% H_2_ in the headspace. Also here, pyrite was not formed (Table S2) while methane production was stimulated more than 10-fold by the added H_2_ (>10 mmol per L of culture). This clearly showed that methanogenesis could be uncoupled from pyrite formation and is essential for the latter, most likely to keep H_2_ or another electron carrier at a low level, to make pyrite formation energetically more favorable as is typically observed in syntrophic processes (26). Although we could not identify the exact electron carrier, molecular H_2_ is the most likely candidate because it was previously observed in abiotic experiments of pyrite formation from FeS and H_2_S (3, 27). Irrespective of the actual electron carrier, syntrophic coupling of pyrite formation to methanogenesis would be exergonic and result in an expected ratio of formed pyrite to methane of 4:1 (equ. 5).

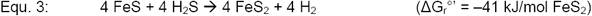

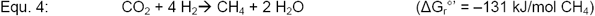

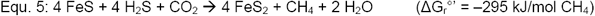

Indeed, the ratio of pyrite to methane formed in culture J5 was 4.1:1 and 3.2:1 in two independent experiments at 28°C (Table S2), respectively, thus confirming the proposed overall reaction stoichiometry.

### Exergonic pyrite formation supports microbial growth

Culture J5 was transferred more than 20 times on medium containing FeS, H_2_S and CO_2_ as sole substrates. This indicates a strict dependence on syntrophic pyrite formation as the only energy-yielding reaction observed. Within incubation periods of 83–248 days, cell densities increased by more than one order of magnitude, from 2×10^5^ cells mL^−1^ to 2–9×10^6^ cells mL^−1^ (Fig. S3). Cells were typically found to be attached to mineral surfaces (Fig. 4*C*); however, there was no indication of cell encrustation (Fig. 4*D*). Assuming an average cell volume of about 1 μm^3^ and a dry mass content of 33% (28), our measured cell densities correspond to a maximum of ca. 3 mg dry cell mass L^−1^. If formation of ATP requires 60–70 kJ mol^−1^ (29) and if – under ideal growth conditions – 10.5 g biomass (dry weight) can be synthesized at the expense of 1 mol ATP (30), a complete conversion of 5 mM FeS + 5 mM H_2_S according to equ. 5 could yield 4–5 mM ATP or ca. 50 mg dry cell mass L^−1^. Of course, the conditions of lithoautotrophic growth in our enrichment cultures are entirely different from those used in the growth experiment by Bauchop and Elsden, with heterotrophic growth in an organically rich medium. Moreover, formation of pyrite from FeS is an extremely slow process catalyzed at or close to mineral surfaces with very slow substrate turnover, which implies that major amounts of metabolic energy have to be invested in cell maintenance without concomitant growth (31). Thus, it is not surprising that our cell yields are lower than estimated above. If every partial reaction in equ. 5 (5 in total; 4 × FeS transformation, 1 × CH_4_ formation) uses about 20 kJ for synthesis of a minimum of 1/3 of an ATP equivalent (29), the total process could synthesize 5/3 mmol ATP equivalents per liter. Considering that in culture J5 a maximum of 62.5% FeS conversion to FeS_2_ was observed (Table S2), the expected cell yield would – optimally – be 12 mg cell dry matter L^−1^, which is close to the measured cell yield.

**Figure 4.**
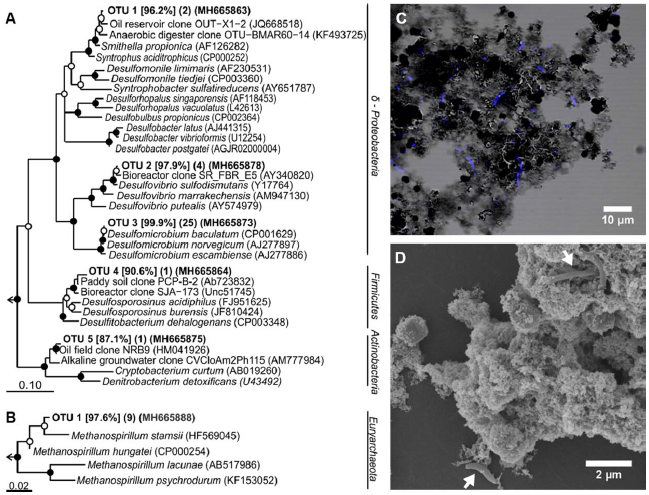
Bacterial and archaeal community composition of enrichment culture J5. RAxML trees based on bacterial (A) and archaeal (B) 16S rRNA gene sequences obtained from culture J5. Representative sequences of OTUs at the approximate species level (99% sequence identity) are shown. Sequence identity to the next cultured relative is given in percent in square brackets. Numbers of clones from the same OTU are presented in parenthesis followed by the GenBank accession number of a representative sequence. Bootstrap support is indicated by closed (≥90%) and open (≥70%) circles at the respective branching points. The scale bar represents 10% (Bacteria) and 2% (Archaea) estimated sequence divergence. (C) Combined phase contrast image and fluorescent image of DAPI-stained cells and (D) scanning electron microscopy image with cells indicated by white arrows of culture J5 after 7.4 and 10 months of incubation, respectively.

The community composition of culture J5 was analyzed by 16S rRNA gene clone libraries. All 16S rRNA gene sequences derived from amplification with a universal archaeal primer set belonged to the same species-level operational taxonomic unit (OTU, 99% sequence identity) and showed 97.6% sequence identity to *Methanospirillum stamsii* (Fig. 4*B*), a hydrogenotrophic methanogen isolated from a low-temperature bioreactor (32). Methanogenesis in culture J5 is likely performed by this OTU. Using a universal bacterial primer set, we detected five bacterial OTUs (Fig. 4*A*). The majority of clones (76%) belonged to OTU 3, which showed 99.9% sequence identity with *Desulfomicrobium baculatum*, a sulfate reducer within the class Deltaproteobacteria (33, 34). Further OTUs related to Deltaproteobacteria were OTU 2 (12% rel. abundance) and OTU1 (6% rel. abundance). OTU 2 was closely related to *Desulfovibrio sulfodismutans* (97.9% sequence identity), which is capable of dismutating thiosulfate or sulfite (35), while OTU 1 was distantly related (96.2% sequence identity) to *Smithella propionica,* which is known to degrade propionate in syntrophy with methanogens (36). The remaining two bacterial OTUs were either distantly related (<91% sequence identity) to cultured members of the Firmicutes (OTU 4, 3% rel. abundance) or Actinobacteria (OTU 5, 3% rel. abundance). Interestingly, all OTUs belonging to the Deltaproteobacteria and Firmicutes fell into larger clusters that include cultured representatives with a sulfur-related energy metabolism. Therefore, it is tempting to speculate that enzymes operating in the respective sulfur transformation pathways might be involved in the microbial conversion of FeS to FeS_2_.

## Conclusion

Pyrite is produced in massive quantities in today’s sediments (2). However, its formation in nature is far from understood, especially because its nucleation is kinetically hindered (1). The presence of sulfide-producing microorganisms as passive pyrite nucleation sites indicated support for abiotic pyrite formation (13-15), but could not be reproduced in every bacterial model system (6). We show that pyrite formation can be mediated by microorganisms to overcome the kinetic hurdle of nucleation, but as an essential part of their energy metabolism and not just as a mere abiotic side reaction on their cell surface. This may help to explain the ambiguous results published so far on the role of microorganisms in this process. Since we found only one archaeal species closely related to methanogens in our enrichments, it is likely that one or several of the bacterial partners actually catalyze FeS transformation to pyrite. An exciting question currently remaining is whether these bacteria utilize internally the H_2_S pathway (equ. 2) or a combination of sulfide oxidation to zero-valent sulfur coupled to the polysulfide pathway (equ. 1) to finally produce pyrite (Fig. S4).

Our results further showed that the reducing equivalents released from FeS transformation to pyrite can be transferred to methanogenesis. This opens an interesting perspective on the metabolic versatility sustaining the vast deep biosphere inhabiting the Earth’s subsurface (37). While recalcitrant organic matter or H_2_ released by radiolytic cleavage of water (38, 39) have been proposed to sustain the enigmatic life forms of the deep biosphere, there is also experimental evidence of a cryptic sulfur cycle within deep sediments that would include pyrite formation coupled to methanogenesis to be functional (17). Our results show that this missing link could indeed be mediated by microorganisms and supply energy to support microbial growth. Since the redox potential (E_h_^°’^) of the FeS/FeS_2_ couple is –620 mV at circumneutral pH (40), it is well suited to provide reducing equivalents also for CO_2_ fixation to acetate (E_h_^°’^ = –290 mV) and more complex organic matter in the pyrite-forming microorganism. Wächtershäuser proposed in his “iron-sulfur world” theory that exactly this mechanism was the basis for an autocatalytic metabolism and the resulting evolution of life at hydrothermal vents on primordial Earth (e.g. 20, 41). Our enrichment cultures may serve as a model to understand the enzymatic background of this hypothesis.

## Methods

### Cultivation

Enrichment cultures were initiated and maintained in carbonate-buffered, sulfide-reduced (1 mM) freshwater mineral medium (42) supplemented with selenite-tungstate solution (43), 7-vitamin solution (42), and trace element solution SL10 (44). The medium was prepared and stored under a N_2_/CO_2_ atmosphere (80:20). The final pH was adjusted to 7.2 to 7.4. From a CO_2_-neutralized sulfide stock solution (45), 350 µmol H_2_S was added to 70 mL mineral medium in 180 mL serum bottles that were sealed with butyl rubber stoppers. Since 1 mM H_2_S was already present as reducing agent in the mineral medium, the final amount of H_2_S was 420 µmol. FeS was prepared from anoxic solutions of 0.4 M FeCl_2_ and 0.4 M Na_2_S. The resulting FeS precipitate was washed at least once and re-suspended in oxygen-free distilled water. For Mössbauer spectroscopy analysis, FeS was prepared from a FeCl_2_ solution that contained 10% ^57^FeCl_2_ to enhance signal quality. 350 µmol FeS was added to 70 mL mineral medium. Enrichment cultures were incubated in the dark at 28°C if not indicated otherwise. For inhibition experiments, cultures were supplemented with either penicillin-G (1,000 U mL^−1^) or 2-bromoethanesulfonate (10 mM). Abiotic controls were run without inoculum.

### Monitoring of substrate turnover

For total dissolved sulfide measurements [Σ(H_2_S_aq_, HS^−^, S^2–^)], 100-µL samples were taken from liquid cultures without disturbing the precipitated iron sulfide minerals, and directly transferred to 100 µL of an anoxic 0.2 M NaOH solution. From the alkalinized sample, 10– 20 µL were fixed in 100 µL of a 0.1 M zinc acetate solution, and sulfide was quantified by the methylene blue method (46). The corresponding amount of H_2_S in the headspace was calculated using Henry’s law and a temperature-adjusted k-value of 0.093 (28°C, 47). CH_4_ was measured by gas chromatography with a flame ionization detector (SRI Instrument SGI 8610C) using a consecutive arrangement of a Porapak (80/100 mesh; 1 m × 2 mm) and a Hayesep-D packed column (80/100 mesh; 3 m × 2 mm). The injector and column temperatures were 60°C, and the detector temperature was 135°C. The carrier gas was N_2_ at a flow rate of 3.2 ml min^−1^. Chromatograms were recorded with the PeakSimple v4.44 chromatography software.

Iron-sulfide minerals were analyzed by ^57^Fe Mössbauer spectroscopy. Within an anoxic glovebox (100% N_2_), the enrichment culture was passed through a 0.44-µm filter and then sealed between two pieces of air-tight Kapton tape. Samples were transferred to a Mössbauer spectrometer (WissEl, Starnberg) within an airtight bottle filled with 100% N_2_ that was only opened immediately prior to loading the samples inside the closed-cycle exchange gas cryostat (Janis cryogenics). Measurements were collected at a temperature of 5 K with a constant acceleration drive system (WissEL) in transmission mode with a ^57^Co/Rh source and calibrated against a 7 µm thick α-^57^Fe foil measured at room temperature. All spectra were analyzed using Recoil (University of Ottawa) by applying the Voight Based Fitting (VBF) routine (48). The half width at half maximum (HWHM) was fixed to a value of 0.138 mm s^−1^ for all samples.

X-ray diffraction (XRD) patterns were recorded with the D8 Discover system (Bruker) with IµS radiation source (2 mm in diameter), and a Lynxexe XE detector. Samples were dried for 2 h under a continuous stream of 100% N_2_ and measured within 48 h under air as described previously (49). Measurements were done using CuKα rays in angles ranging from 10–70° 2θ in 0.02° steps with 2,880 sec measuring time and a total measuring time of 12 h and 47 min. The resulting spectra were compared with spectra provided in the international crystal structure database (ICSD), FIZ Karlsruhe (version 2016/2) using the software DIFFRAC.EVA (version 4.1.1, Bruker).

### Scanning electron microscopy coupled to energy dispersive x-ray spectroscopy (SEM-EDX)

For SEM-EDX analysis, 1 mL of culture was centrifuged at 4,500 ×*g* for 10 min. 200 µL of the resulting pellet was transferred on gelatin-coated glass slides. Samples were fixed in 1 mL 2.5% glutaraldehyde in 0.1 M HEPES-buffer containing 0.01 M KCl (HEPES-KCl) and in 2% OsO_4_ in HEPES-KCl for 60 min each. Fixed samples were dehydrated in a graded ethanol series (30%, 50%, 70%, 80%, 90%, 96% and absolute ethanol) for 30 min each. Thereafter, samples were critical-point dried under CO_2_ in a Bal-Tec CPD030 (Balzers). Sputter coating of 6 nm platinum was done in a Quorum Q150R ES sputter coater (Quorum Technologies) and micrographs were taken with a FESEM Auriga 40 (Zeiss). EDX mappings and point measurements were taken at a working distance of 5 mm with an Oxford X-Max detector (Oxford Instruments) and at 10 kV and 15 kV, respectively. Point measurements were normalized to 10,000 counts within a Kα energy of 6.3–6.5 keV. Sample preparation for cell counts by fluorescent microscopy is described in detail in the Supporting Information.

### Phylogenetic analysis

Total genomic DNA was extracted from 50 mL of a 4.5-months old culture using a phenol-based beat-beating protocol modified after Loy, Beisker and Meier (50). Subsequent amplification of bacterial or archaeal 16S rRNA genes was done using standard PCR protocols based on universal primers. Details are given in the Supporting Information. 16S rRNA gene clone libraries were constructed using the TOPO^®^ TA Cloning^®^ Kit (ThermoFisher Scientific). Bacterial or archaeal 16S rRNA gene fragments were aligned by use of the SINA webaligner (51) to the non-redundant 16S rRNA gene database v.123.1 available on the SILVA online platform (52, www.arb-silva.de) and imported into ARB for initial phylogenetic analysis (53). OTU clustering was performed in mothur v.1.22.2 (54) using the furthest neighbor approach and a 99%-identity cutoff to delineate OTUs at the approximate species level (55). For phylogenetic inference of 16S rRNA gene fragments representing individual OTUs, Maximum Likelihood (ML) trees were calculated using RAxML v8.2.9 (56) as implemented on the CIPRES webserver (57, www.phylo.org). Using a 50% conservation filter of nucleic acid positions within the domain Bacteria, a RAxML tree was inferred from 1,102 unambiguously aligned nucleic acid positions for bacterial 16S rRNA genes. The reconstruction of the archaeal tree followed the same outline but using 752 unambiguously aligned nucleic acid positions and no conservation filter because of the close relatedness of all included sequences. Calculations were based on the GTRGAMMA distribution model of substitution rate heterogeneity. MRE-based bootstrap analysis stopped after 204 and 102 replicates for the bacterial and archaeal 16S rRNA gene tree, respectively. Sequences are available from NCBI GenBank under accession numbers MH665848–MH665880 and MH665881–MH665889 for Bacteria and Archaea, respectively.

## Acknowledgements

This study was financed by the Konstanz Research School Chemical Biology (KoRs-CB). We are grateful to Waltraud Dilling for maintaining initial pyrite-forming enrichments and to Ben Griffin for his initial work on the latter. We further thank Michael Laumann for his engagement with SEM-EDX, Stephan Siroky and the Particle Analysis Center for the XRD analysis, as well as the Bioimaging Center (University of Konstanz).

## Supporting Information

### Supporting Materials and Methods

#### Cell counts by fluorescence microscopy

For cell counts, 0.5 mL culture was fixed overnight in 9.5 mL freshly prepared paraformaldehyde solution (4%), subsequently centrifuged at 10,000 × *g* for 10 min at 4°C and re-suspended in 1 mL PBS [130 mM NaCl, 5% (v/v) phosphate buffer (40 mM NaH_2_PO_4_, 160 mM Na_2_HPO_4_), pH 7.2] and 9 mL ammonium oxalate solution (5.6 g ammonium-oxalate and 4.2 g oxalic acid dihydrate in 200 mL distilled water). Samples were vortexed for 10 min to dissolve most of the iron sulfide minerals so that cells could be collected on a 0.2 µm pore size filter (GTTP-white, Millipore). Filters were air-dried and stored at –20°C. Filter sections were stained with a 1 µg mL^−1^ 4’,6-diamidino-2-phenylindole (DAPI) solution and incubated for 10 min in the dark. Thereafter, filters were washed for 5 min in distilled water, followed by two 1-min washing steps in 80% ethanol. Dry DAPI-stained filters were mounted on microscope slides using CitiFluor™ AF1. For fluorescence microscopy, an inverted microscope (AxioObserver, Zeiss) with a 40x/0.60 LD-PlanNeofluar objective was used. Z-stacks were acquired with a distance of 0.28 µm. Image processing involved 3-dimensional deconvolution of each stack using a theoretical PSF with ZEN Black (Zeiss AG). Cells were counted using an image processing workflow set up in KNIME 3.4.0 (58) using orthogonal projections of the de-convoluted input stacks. The workflow is available at https://github.com/bic-kn/cell-counting-workflow.

#### DNA extraction

The DNA extraction protocol was adopted from (50). A 4.5-month old 50-mL culture was harvested after CH_4_ concentrations reached a plateau of 2.1% in the headspace (corresponding to 55 µmol produced CH_4_). Harvesting was done by 10 min of centrifugation at 6,000 × *g*. The pellet was re-suspended in 400 μL autoclaved TE-Buffer (10 mM Tris & 1 mM EDTA in MQ water, pH 8) and stored for three hours at –20°C. Cells were thawed on ice, mixed with heat-sterilized zirconium beads (0.1 mm), 600 μL phenol/chloroform/isoamylalcohol (25:24:1, Carl Roth), and 150 µL of a 10% sterile-filtered SDS-solution in a screw-cap tube, and vigorously shaken for 20 min using a vortexer. After centrifugation for 20 min at 20,817 × *g* and 4°C, the aqueous supernatant was transferred to a new reaction tube. Because the aqueous phase was hardly visible due to remaining iron sulfide minerals, another 10-minute centrifugation step was used to remove residual phenol from the extract. DNA was precipitated by incubation at –20°C overnight in 0.1 volume of 3 M Na-acetate (in MQ-water, autoclaved) and 2.5 volumes absolute ethanol. Afterwards, the pellet was washed twice with 70% ethanol, dried for 5 min, and re-suspended in 50 μL DNase- and RNAse-free H_2_O. DNA concentrations were quantified fluorimetrically using Quant-iT PicoGreen (Invitrogen).

#### 16S rRNA gene clone library

Amplification of bacterial 16S rRNA genes was performed with Bact8f (5’-AGA GTT TGA TYM TGG CTC-3’) as forward primer (59) and 1492r (5’-N TAC CTT GTT ACG ACT-3’) as reverse primer (60). Archaeal species were targeted by AR109F (5’-ACK GCT CAG TAA CAC GT-3’) as forward (61) and AR915 (3’-GTG CTC CCC CGC CAA TTC CT-3’) as reverse primer (62). The PCR mixture contained 0.2 mM of each dNTP, 2 mM MgCL_2_, 20 µg BSA, 1 U of *Taq* DNA polymerase, and a Taq polymerase buffer with KCl (ThermoFisher Scientific). The PCR was performed using an initial denaturation at 95°C for 5 min; 30 cycles of 95°C for 30 s, 50°C for 30 s, 72°C for 1.5 min; and a final elongation at 72°C for 7 min. For PCRs with archaeal primers, the annealing temperature was set to 55°C. Amplification products were purified by use of the Zymo Research DNA Clean & Concentrator Kit (Zymo Research). 16S rRNA clone libraries were obtained with the TOPO^®^ TA Cloning^®^ Kit (ThermoFisher Scientific). Clones were screened by M13-PCR for inserts of the correct size according to the manufacturer’s instructions. Resulting PCR products of expected length were purified by use of the Zymo Research DNA Clean & Concentrator Kit and sent for sequencing.

## Supporting Tables

**Table S1.** Overview of inocula used to establish initial pyrite-forming enrichment cultures.

| Enrichment | Date | Sampling area | Material | Medium† | Temp. (°C) | pH |
| --- | --- | --- | --- | --- | --- | --- |
| J5* | Sept. 1995 | Sewage treatment plant<br>Konstanz, Germany | digested sewage sludge | freshwater | 28 | 7.2 |
| J2* | Sept. 1995 | Sewage treatment plant<br>Konstanz, Germany | digested sewage sludge | freshwater | 16 | 7.2 |
| J7 | Apr. 1993 | Rio Tentor (Venice, Italy) | brackish sediment | marine | 16 | 7.2 |
| J8* | Mar. 1991 | Wadden sea sediment,<br>Groningen, The Netherlands | marine sediment | marine | 16 | 7.2 |
| J9* | Apr. 1993 | Fish market channel<br>(Venice, Italy) | brackish sediment | marine | 16 | 7.2 |
| X1 | Sept. 1995 | Sewage treatment plant<br>Tübingen-Lustnau, Germany | digested sewage sludge | freshwater | 28 | 7.2 |
| X2 | Sept. 1995 | Lake Constance, Güll | freshwater sediment | freshwater | 28 | 7.2 |

† after Widdel and Pfennig (42)
\* CH<sub>4</sub> formation observed

**Table S2.** Iron mineral analysis by Mössbauer spectroscopy at a temperature of 5 K in nearly 7-month-old (207 days) culture J5 incubated at various temperatures and in the presence of various inhibitors. Mössbauer parameters were obtained through Voigt based fitting (VBF). δ – isomer shift, ΔE_Q_ – quadrupole splitting, ε – quadrupole shift, B_hf_ – internal magnetic field, R.A. – relative area, X^2^ – goodness of fit parameter. The absolute amount of formed FeS_2_ was inferred from the relative area of the FeS_2_ signal and the maximum amount of 350 µmol that could be produced if all FeS would have been converted to FeS_2_.

| Sample | Phase | $\delta$<br>(mm/s) | $\Delta E_Q$<br>(mm/s) | $\varepsilon$<br>(mm/s) | $B_{hf}$<br>(T) | R. A.<br>% | $X^2$ | $FeS_2$<br>($\mu mol$) |
| --- | --- | --- | --- | --- | --- | --- | --- | --- |
| Abiotic control | $FeS$ | 0.43 | | 0.15 | 13.1 | 64.3 | 1.67 | |
| | $FeS_x$ | 0.50 | 0.13 | | | 35.7 | | |
| 4 °C | $FeS$ | 0.47 | 0.27 | | | 37.8 | 6.01 | |
| | $FeS_x$ | 0.43 | | 0.07 | 17.2 | 62.2 | | |
| 16 °C | $FeS_2$ | 0.41 | 0.38 | | | 18.5 | 0.78 | 64.8 |
| | $FeS_x$ | 0.39 | | 0.02 | 19.2 | 48.8 | | |
| | $FeS_x$ | 0.42 | | -0.02 | 16.5 | 32.7 | | |
| 28 °C<br>Replicate I | $FeS_2$ | 0.41 | 0.50 | | | 52.8 | 0.88 | 184.8† |
| | $FeS_x$ | 0.32 | | 0.07 | 15.7 | 38.9 | | |
| | $Fe_3S_4$ | 0.46 | | 0.05 | 32.0 | 8.3 | | |
| 28 °C<br>Replicate II | $FeS_2$ | 0.40 | 0.58 | | | 62.5 | 0.78 | 218.8§ |
| | $FeS_x$ | 0.29 | | -0.2 | 16.4 | 31.0 | | |
| | $Fe_3S_4$ | 0.60 | | -0.05 | 32.0 | 6.5 | | |
| 46 °C | $FeS_2$ | 0.42 | 0.41 | | | 39.4 | 1.67 | 137.9 |
| | $FeS_x$ | 0.10 | | -0.09 | 21.1 | 50.5 | | |
| | $FeS_x$ | 0.40 | | 0.05 | 15.7 | 7.8 | | |
| | $Fe_3S_4$ | 0.64 | | 0.03 | 33.2 | 2.3 | | |
| 60 °C | $FeS$ | 0.54 | 0.16 | | | 27.1 | 1.74 | |
| | $FeS_x$ | 0.34 | | 0.05 | 2.4 | 21.6 | | |
| | $FeS_x$ | 0.55 | | -0.08 | 16.6 | 43.9 | | |
| | $Fe_3S_4$ | 0.63 | | -0.08 | 31.9 | 7.3 | | |
| Penicillin + 79% $H_2$<br>( headspace) | $FeS$ | 0.51 | 0.16 | | | 31.6 | 1.12 | |
| | $FeS_x$ | 0.38 | | 0.15 | 5.3 | 17.4 | | |
| | $FeS_x$ | 0.85 | | 0.01 | 23.0 | 37.5 | | |
| | $FeS_x$ | 0.46 | | -0.07 | 15.5 | 13.4 | | |
| Penicillin | $FeS$ | 0.46 | 0.27 | | | 26.4 | 1.15 | |
| | $Fe_3S_4$ | 0.68 | | -0.04 | 33.1 | 8.8 | | |
| | $Fe_3S_4$ | 0.52 | | 0.05 | 31.7 | 18.8 | | |
| | $FeS_x$ | 0.44 | | 0.00 | 14.9 | 15.0 | | |
| | $FeS_x$ | 0.43 | | 0.02 | 20.2 | 31.1 | | |
| BES | $FeS$ | 0.50 | 0.22 | | | 28.1 | 0.76 | |
| | $Fe_3S_4$ | 0.69 | | -0.15 | 29.6 | 23.4 | | |
| | $Fe_3S_4$ | 0.62 | | 0.02 | 33.1 | 7.1 | | |
| | $FeS_x$ | 0.25 | | 0.15 | 7.4 | 20.2 | | |
| | $FeS_x$ | 0.44 | | -0.16 | 16.6 | 21.2 | | |

† corresponding amount of formed $CH_4$ : 44.9 $\mu mol$
§ corresponding amount of formed $CH_4$ : 67.6 $\mu mol$

## Supporting Figures

**Figure S1.**
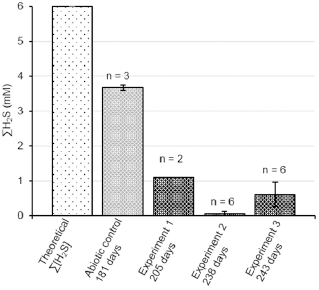
Total H_2_S as the sum of H_2_S_gaseous_, H_2_S_aqueous_, HS^−^, and S^2–^ in the non-inoculated medium as compared to the abiotic control and enrichment culture J5 in various independent incubation experiments. The time of incubation is indicated in days. Biological replicates are indicated as n.

**Figure S2.**
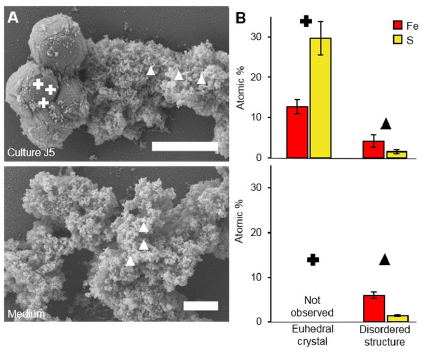
Fe:S ratio of different mineral phases in culture J5 and freshly prepared medium without inoculum. (A) Exemplary scanning electron microscopy images used as guidance to perform energy dispersive X-ray spectroscopy (EDX) point measurements of culture J5 after nearly 10 months of incubation (295 days) and of freshly prepared medium without inoculum. Scale bars represent 2 µm. Symbols in the SEM images indicate EDX point measurements (crosses for crystals, triangles for disordered structure). (B) Atom percent ratio of iron (red) and sulfur (yellow) as derived from EDX point measurements of euhedral crystals resembling pyrite as well as disordered structures resembling the sum of the remaining Fe-S-mineral phase. Measurements were done on eight different sampling areas with three EDX point measurements each.

**Figure S3.**
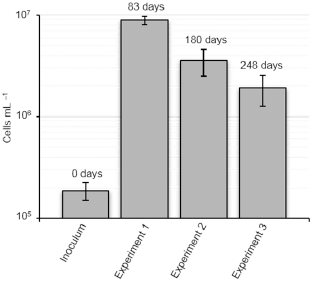
Average cell counts of culture J5 as based on DAPI-stained cells in three independent incubation experiments at 28°C as compared to freshly inoculated medium. The time of incubation is indicated in days. Data was obtained from biological duplicates, each measured in technical triplicates. Standard deviations are given for technical replicates.

**Figure S4.**
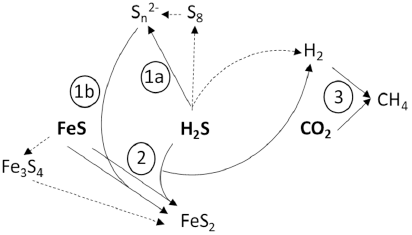
Schematic overview of potential FeS_2_ formation pathways in culture J5. The scheme illustrates pyrite formation either by a combination of sulfide oxidation to zero-valent sulfur (1a) coupled to the polysulfide pathway (1b) or by the H_2_S pathway (2). The released reducing equivalents are likely transferred in the form of H_2_ to methanogenesis (3) to reduce CO_2_ to CH_4_. Dashed lines leading to *cyclo*-octasulfur (S_8_) and greigite (Fe_3_S_4_) represent potential alternative pathways or side reactions, respectively.

